# Temporal regularity and stimulus control in multiple fixed interval schedules

**DOI:** 10.1101/695353

**Authors:** Estela B. Nepomoceno, André M. Cravo, Marcelo B. Reyes, Marcelo S. Caetano

**Author notes:** Corresponding author at Center for Mathematics, Computation, and Cognition, Federal University of ABC (UFABC), Alameda da Universidade s/n, São Bernardo do Campo, SP, Brazil. CEP: 09606-045. *E-mail address* (M. S. Caetano).

## Abstract

In multiple fixed interval schedules of reinforcement, different time intervals are signaled by different environmental stimuli which acquire control over behavior. Previous work has shown that temporal performance is controlled not only by external stimuli but also by temporal aspects of the task, depending on the order in which the different intervals are trained – intermixed across trials or in blocks of several trials. The aim of this study was to further describe the training conditions under which the stimuli acquire control over temporal performance. We manipulated the number of consecutive trials of each fixed interval (FI) per training block (Experiment I) and the number of FIs trained (Experiment II). The results suggest that when trained in blocks of several consecutive trials of the same FI, temporal performance is controlled by temporal regularities across trials and not by the visual stimuli that signal the FIs. One possible account for those data is that the temporal cues overshadow the visual stimuli for the control of temporal performance. Similar results have also been observed with humans, which suggest that temporal regularity overcomes the stimuli in the control of behavior in temporal tasks across species.

## 1. Introduction

In fixed-interval (FI) schedules of reinforcement, the first operant (target) response after a period of time has elapsed is rewarded, while earlier responses have no effect (Catania, 1999; Ferster & Skinner, 1957). After some sessions of training with FI schedules, little to no responses are observed at the beginning of the interval when food availability is distant, and as the interval elapses, response rate increases (Ferster & Skinner, 1957; Pavlov, 1927). This pattern of responding is termed *temporal gradient* and the development of the difference in response rate early vs. later in the interval is used as evidence of learned *temporal discrimination* (Skinner, 1938).

In multiple FI schedules, different stimuli signal different intervals. After some sessions of training with multiple FI schedules, the increase in response rate within-trials seems to be guided by the stimulus presented in each trial. Response rate rises earlier in trials signaled by stimuli which have been paired with shorter intervals, while response rate rises later in trials with stimuli signaling longer intervals. This discrimination across FIs is called *discrimination of interval* (Ferster & Skinner, 1957).

It is assumed that discrimination of interval is a result of an association between the different stimuli and their respective signaled intervals, developed during training. However, other strategies may lead to interval and temporal discriminations that do not require associations between stimuli and FIs to be formed (Marshall & Kirkpatrick, 2015). Specifically, when trained in blocks of several consecutive trials of the same signaled FI, the temporal regularity across trials seems to control temporal performance, as opposed to the discriminative stimuli – even though these stimuli are still reliable cues to the FI.

These counterintuitive results have been reported for both rats (Caetano, Guilhardi, & Church, 2007, 2012) and humans (Guilhardi, Menez, Caetano, & Church, 2010; Labliuk, Guilhardi, Cravo, Church, & Caetano, 2015). Caetano and cols. (2007, 2012), for example, showed that rats trained with three signaled FIs in trials presented intermixed in the same session respond differently to each signaling stimulus (i.e., rats seem to associate the stimuli with their respective intervals), but not when trained in consecutive blocks of 10 sessions with the same FI, or in alternating blocks of 1 session per FI. Both in the intermixed and blocked designs, the number of overall trials of each signaled FI is identical; the only difference is the order in which they were trained.

In humans, Guilhardi and colleagues (2010) trained multiple FIs with a modified peak procedure task presented as a shooting game. The goal was to hit a moving target when it crossed the center of the screen. This moment could be predicted by a visual stimulus (background color) or by the temporal regularity in the task (same time as in previous trials). Similar to the results observed in rats, participants used the temporal regularity in the task, instead of the visual stimuli, to predict when to shoot.

In the present study, we further describe the conditions under which stimulus or temporal control of performance is observed in rats. Experiment I assessed the role of the number of consecutive trials within-blocks in the emergence of stimulus or temporal control of performance; and Experiment II investigated whether training a smaller number of signaled FIs would lead to stimulus control.

## 2. Experiment I

Experiment I tested how the temporal regularity of the task - gauged by the number of consecutive trials of the same FI – modulates the temporal performance of rats. Different rats were trained with consecutive blocks of 5, 20, or 60 trials of each FI, and by the end of training the number of trained trials of each FI were identical across groups. The working hypothesis was that, as block size decreases, so does the temporal regularity, leading to a stimulus-controlled performance.

### 2.1 Method

All experimental procedures were approved by the Animal Care and Use Committee at Universidade Federal do ABC (UFABC) and conform to guidelines for the Ethical Treatment of Animals (National Institutes of Health).

#### 2.1.1 Animals

Eighteen male Wistar rats (purchased from Federal University of São Paulo, São Paulo, SP) arrived in the lab at 30 days of age. They were housed in a colony room with a 12-hr light:dark cycle (lights on at 7 a.m.) in groups of three rats per home cage. All rats were handled daily for 111 days before the beginning of training. Water was freely available in the home cages. Daily ration consisted of up to 60 pure sucrose pills nº 5 (45 mg) delivered during the experimental sessions, and of an additional 45 g of Nuvilab CR-1 food pellets delivered in the home cages after the training sessions (i.e., 15 g per rat, on average). Training began when they were 144 days old and occurred in the morning and afternoon shifts.

#### 2.1.2 Apparatus

Six standard Med Associates Inc. operant chambers, 25-cm wide, 30-cm high, by 30-cm deep, were used during training. Each chamber was equipped with a magazine pellet dispenser, which delivered sucrose pellets into a food cup. The food cup was mounted on the front wall of the chamber. Head-entry responses into the food cup were detected by a pair of LED photocells. Three visual stimuli were used during training and provided diffuse illumination of approximately 200 Lux. They were (a) the light located above the left lever, (b) the houselight, and (c) all three lights flickering (0.5-s on/0.5-s off) simultaneously. Each chamber also had two fixed levers, which were not used in the present experiment. A Celeron D/504 computer running Med-PC IV for Windows XP 2002 controlled and recorded all experimental events with 2-ms resolution.

#### 2.1.3 Procedure

Rats were trained on a multiple-fixed-interval procedure with three intervals (30 s, 60 s or 120 s) signaled by the three different visual stimuli for 85 daily sessions (Monday through Saturday). The assignment of stimuli to intervals was counterbalanced across rats. Experimental sessions consisted of 60 trials or 140 minutes, whichever came first. In each trial, food was primed 30 s, 60 s or 120 s seconds after stimulus onset. The first head entry into the food cup after the food was primed delivered the sucrose pellet, terminated the stimulus, and started a 20-s period with no stimulus (intertrial interval, ITI). Head entries made prior to the time of food prime were recorded but had no effect.

##### 2.1.3.1 Phase 1A – Blocked training with fixed ITI

In the first phase (48 sessions), rats were divided into three groups and trained in a blocked design. For the first group (G60; n = 6), the three signaled FIs were trained in blocks of 60 consecutive trials, i.e., one FI per session. They were selected by a pseudo-random permutation of the three possibilities, with the restriction that the same FI could not be trained in two consecutive experimental sessions. For the second group (G20; n = 6), the FIs were trained in three blocks of 20 consecutive trials, so the three FIs were trained in each session. The training order of intervals in each experimental session was defined by a random permutation of the three possibilities. Finally, for the third group of rats (G5; n = 6), the three FIs were trained in 12 blocks of 5 consecutive trials (4 blocks of each FI), also totaling 60 trials in the session. The order for the 12 blocks was defined by a pseudorandom permutation of the three possibilities, with the restriction that the same FI could not be trained in two consecutive blocks. The ITI between trials for all groups was fixed at 20 seconds.

##### 2.1.3.2 Phase 1B – Blocked training with varied ITI

In phase 1B, beginning in day 49 of training, the ITI changed from a fixed 20 s to a fixed 10 s plus a random 8-s portion (tandem FT10-s + RT8-s) with a maximum random portion of 16 s. Therefore, the total ITI varied from 10 to 26 s. This change was made to control for the possibility that rats were using the interval between food delivery and the next food prime as a cue to the interval trained. Phase 1B was trained for 19 sessions.

##### 2.1.3.3 Phase 2 – Intermixed training

In the second phase (18 sessions), all rats were trained with the three FIs intermixed within each session (20 trials each per session). The FI trained in each set of three trials was defined by a pseudorandom permutation of the three pairs without replacement, with the restriction that the same FI could not be trained in two consecutive trials (i.e., the FI selected for the first trial in each triad could not coincide with the FI trained in the last trial of the previous triad).

#### 2.1.4 Data analysis

Mean head-entry rate as a function of time from the onset of the stimulus (i.e., temporal gradient) was used to describe performance during training, as previously described by Guilhardi and Church (2005). For statistical comparisons of performance in phase 1A with 1B, the last three sessions in phase 1A and first three sessions in phase 1B for each FI were used. For comparisons between blocked and intermixed training, data from the last three sessions of blocked and intermixed training for each FI were used. Then, the sessions-averaged temporal gradients were smoothed individually with a moving window of size 3 seconds to reduce local variability. Finally, mean response rate during the first 30 seconds of each FI was calculated and used in the comparisons. These 30 seconds are the common period of time across the three FIs, and has been previously used to compare performance across different intervals (*response rate at comparable intervals;* Guilhardi and Church, 2005).

For comparisons of performance between phase 1A and 1B, response rates at comparable intervals were tested with repeated-measures ANOVAs. Post hoc Bonferroni corrections were applied where appropriate and partial eta squared or Cohen’s *d* was used to describe effect sizes. Levene’s test of normality or Shapiro-Wilk was used before all inferential analyses, and revealed no deviations from normality. All rates and absolute distances are shown as mean (±SEM).

Three rats (one from each group) did not discriminate across FIs during the intermixed condition and were excluded from the analyses. For these rats, the difference between mean response rates during the 30- and 120-s FIs in all intermixed sessions was smaller than 2.5 SEM of the same difference based on the group average. A visual inspection of the temporal gradients for those three rats confirmed their superposition. Differential responding in intermixed trials was needed to confirm that animals were able to discriminate the stimuli and that any gradient superposition during blocked training would not be due to an inability to visually discriminate between them.

### 2.2 Results

After blocked training, temporal performance did not differ between phases 1A and 1B for any group. Mean rates (and standard errors) for each interval in the last three sessions of phase 1A, and in first three sessions of phase 1B are shown in Table 1. A repeated-measures ANOVA for each group, with interval (30- vs. 60-vs. 120-s FI) and phase (1A vs. 1B) as within-factors, revealed a significant effect of interval for group G60 (*F*_*2,8*_ = 49.51, *p* < 0.001, η^²^_p_ = 0.925), for group G20 (*F*_*2,8*_ = 96.42, *p* < 0.001, η^²^_p_ = 0.960), and for group G5 (*F*_*2,8*_ = 76.34, *p* < 0.001, η^²^_p_ = 0.950). As there was no significant effect of phase nor any interactions, data from phases 1A and 1B were combined (totaling 67 sessions) and all remaining analyses for the blocked condition were performed on the last three sessions.

**Table 1:**
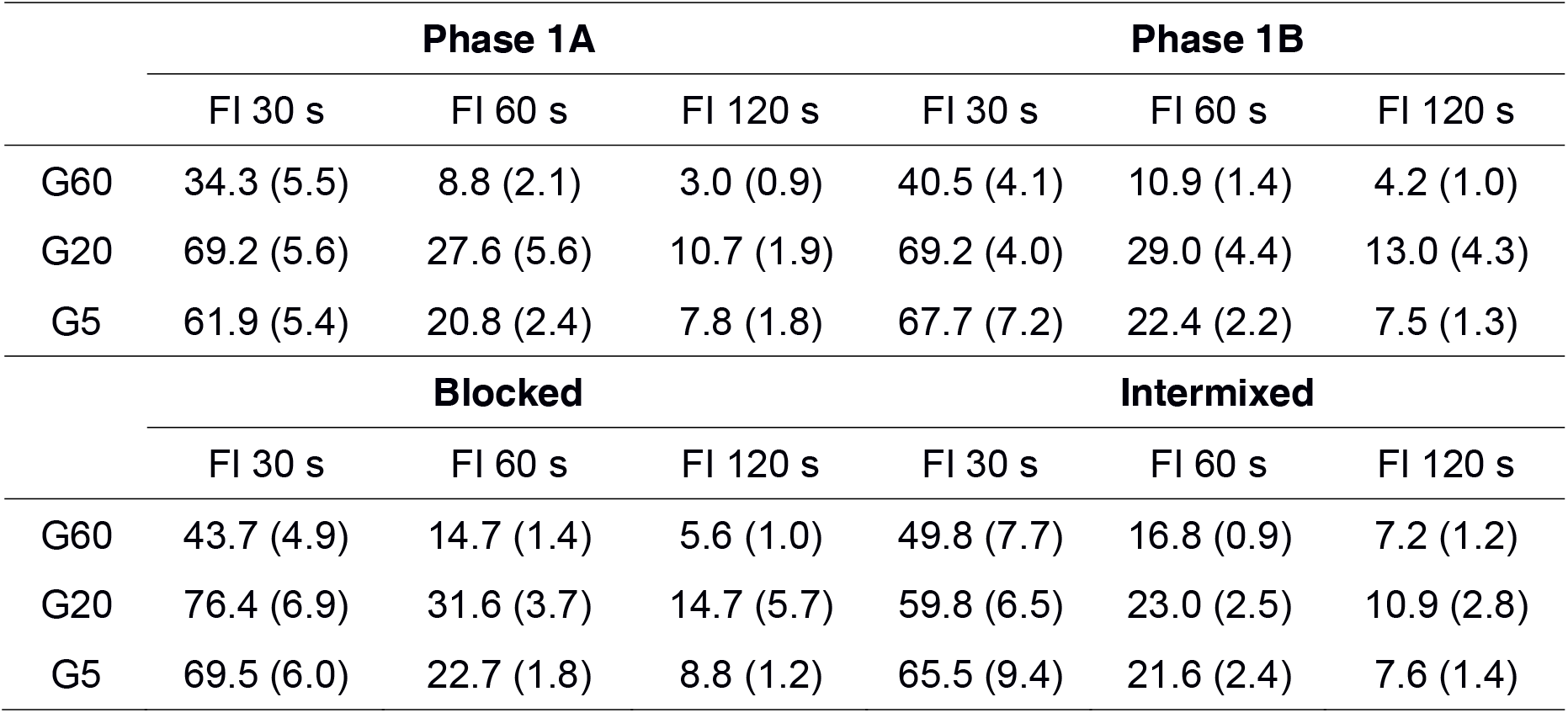
Mean response rate (and standard error) in responses per minute during the first 30 s of each response gradient for all trials in different training phases.

|  | Phase 1A |  |  | Phase 1B |  |  |
| --- | --- | --- | --- | --- | --- | --- |
|  | FI 30 s | FI 60 s | FI 120 s | FI 30 s | FI 60 s | FI 120 s |
| G60 | 34.3 (5.5) | 8.8 (2.1) | 3.0 (0.9) | 40.5 (4.1) | 10.9 (1.4) | 4.2 (1.0) |
| G20 | 69.2 (5.6) | 27.6 (5.6) | 10.7 (1.9) | 69.2 (4.0) | 29.0 (4.4) | 13.0 (4.3) |
| G5 | 61.9 (5.4) | 20.8 (2.4) | 7.8 (1.8) | 67.7 (7.2) | 22.4 (2.2) | 7.5 (1.3) |
|  | Blocked |  |  | Intermixed |  |  |
|  | FI 30 s | FI 60 s | FI 120 s | FI 30 s | FI 60 s | FI 120 s |
| G60 | 43.7 (4.9) | 14.7 (1.4) | 5.6 (1.0) | 49.8 (7.7) | 16.8 (0.9) | 7.2 (1.2) |
| G20 | 76.4 (6.9) | 31.6 (3.7) | 14.7 (5.7) | 59.8 (6.5) | 23.0 (2.5) | 10.9 (2.8) |
| G5 | 69.5 (6.0) | 22.7 (1.8) | 8.8 (1.2) | 65.5 (9.4) | 21.6 (2.4) | 7.6 (1.4) |

Figure 1 shows mean response rate as a function of time from stimulus onset for groups G60 (top row), G20 (middle row), and G5 (bottom row) during blocked (left column) and intermixed training (right column). Temporal gradients were averaged across all trials within the last trained sessions. Rats from all groups similarly discriminated across the three FIs during blocked and intermixed training. A repeated-measures ANOVA, with interval (30- vs. 60-vs. 120-s FIs) and training (blocked vs. intermixed) as within-subject factors, revealed a significant effect of interval for the G60 group (*F*_*2,8*_ = 30.91, *p* < 0.001, η^²^_p_ = 0.885), a significant effect of interval (*F*_*2,8*_ = 92.79, *p* < 0.001, η^²^_p_ = 0.959) and a marginal effect of training (*F*_*1,4*_ = 8.63, *p* = 0.042, η^²^_p_ = 0.683) for the G20 group, and a significant effect of interval for the G5 group (*F*_*2,8*_ = 55.21, *p* < 0.001, η^²^_p_ = 0.932). No interactions were found. Post hoc comparisons between gradients within each panel showed a statistically significant difference between all pairs of gradients in each panel (i.e., between 30- and 60-s, 30- and 120-s, and 60- and 120-s; Table 1). In general, these results suggest that rats were able to discriminate the different intervals similarly in the blocked and intermixed conditions.

**Figure 1.**
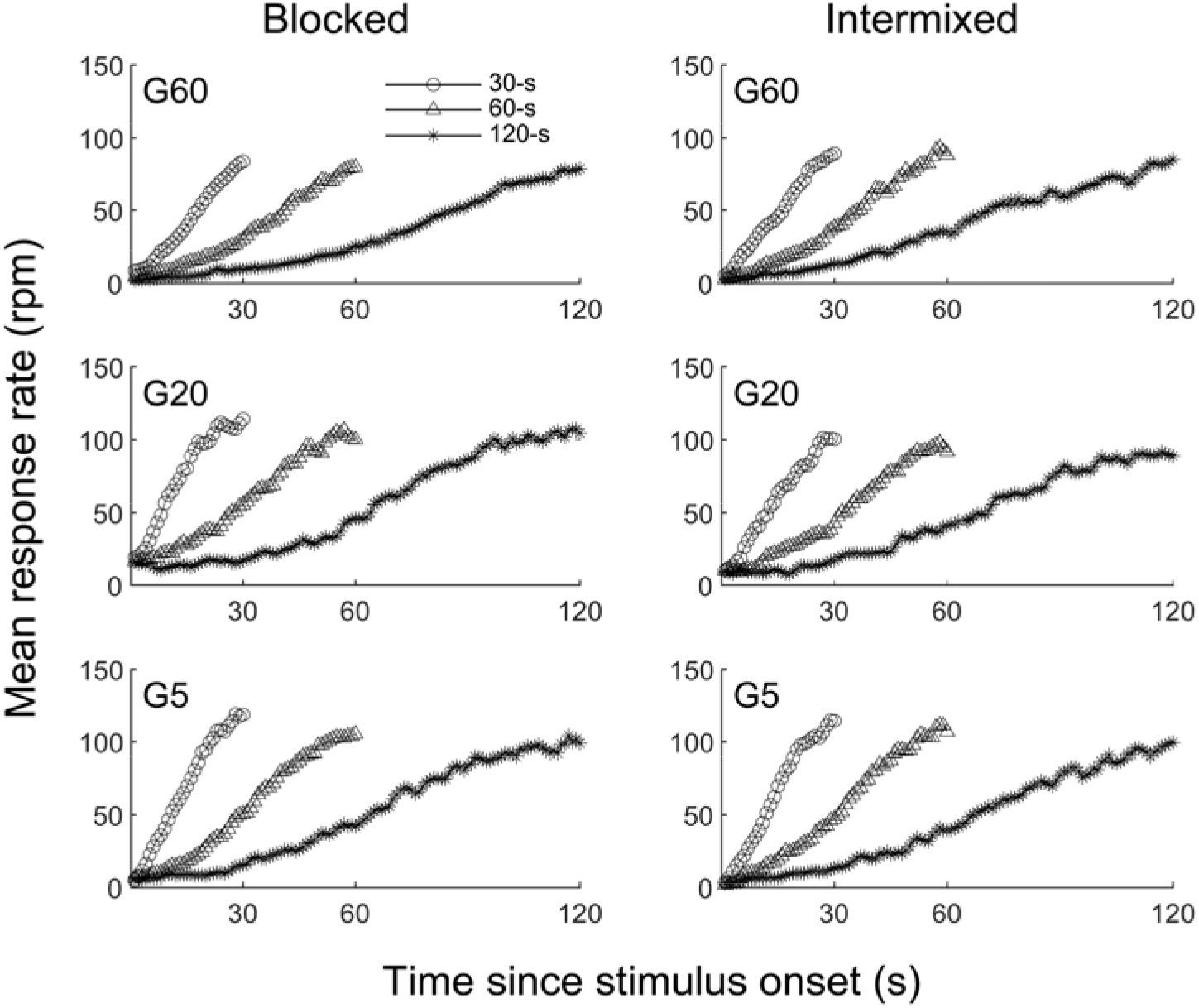
Mean response rate during the FI 30, FI 60, and FI 120 s as a function of time since stimulus onset for all trials in the blocked (left column) and intermixed (right column) conditions for groups G60 (upper panels), G20 (middle panels), and G5 (lower panels).

Although average gradients suggest similar performance at the end of blocked and intermixed training for all groups, a trial-by-trial analysis suggests a within-session adjustment in performance depending on block size (see Table 2 for summary statistics). Figure 2 shows mean response rate as a function of time from stimulus onset for groups G60 (top row), G20 (middle row), and G5 (bottom row) in the first (left column) and last (right column) trials of the last three sessions of blocked training. Interval discrimination in the first trial seems to differ across groups. However, by the last trial within blocks, the gradients were separated for all groups. A repeated-measures ANOVA with interval (30- vs. 60- vs. 120-s FIs) and trial (first vs. last) as within-subject factors revealed a significant effect of interval (*F*_*2,8*_ = 6.26, *p* = 0.023, η^²^_p_= 0.610) and an interval x trial interaction (*F*_*2,8*_= 12.50, *p* = 0.003, η^²^_p_ = 0.758) for the G60 group; a significant effect of interval for the G20 group (*F*_*2,8*_ = 33.76, *p* < 0.001, η^²^_p_ = 0.894); and a significant effect of interval (*F*_*2,8*_ = 47.25, *p* < 0.001, η^²^_p_ = 0.922) and trial (*F*_*1,4*_ = 15.36, *p* = 0.017, η^²^_p_ = 0.793) for the G5 group. The significant interaction in G60 expresses the within-session adjustment in performance for this group. Post hoc comparisons between gradients in the first and last trial for group G60 revealed significant differences across intervals only in the last trial (30 vs. 120 s, *t*_*4*_ = 4.48, *p* = 0.011, Cohen’s *d* = 2.004; 60 vs. 120 s, *t*_*4*_ = 6.99, *p* = 0.002, Cohen’s *d* = 3.126). For group G20, post hoc comparisons revealed a significant difference between 30 and 120 s (*t*_*4*_ = 4.02, *p* = 0.016, Cohen’s *d* = 1.798) in the first trial; and significant differences between 30 and 60 s (*t*_*4*_ = 4.63, *p* = 0.010, Cohen’s *d* = 2.068) and between 30 and 120 s (*t*_*4*_ = 19.23, *p* < 0.001, Cohen’s *d* = 8.599) in the last trial. Finally, for group G5 post hoc comparisons showed a significant difference between 30 and 120 s (*t*_*4*_ = 3.82, *p* = 0.019, Cohen’s *d* = 1.709) in the first trial; and significant differences between 30 and 60 s (*t*_*4*_ = 3.41, *p* = 0.027, Cohen’s *d* = 1.526) and between 30 and 120 s (*t*_*4*_ = 5.67, *p* = 0.005, Cohen’s *d* = 2.534) in the last trial. Overall, these results show that rats in G60 were the only ones which did not discriminate the different FIs in the first trial of the final sessions of training.

**Table 2:** Mean response rate (and standard error) in responses per minute during the first 30 s of each response gradient in first and last trials shown in Figures 2 and 3.

|  |  | **Blocked** |  |  | **Intermixed** |  |  |
| --- | --- | --- | --- | --- | --- | --- | --- |
|  |  | FI 30 s | FI 60 s | FI 120 s | FI 30 s | FI 60 s | FI 120 s |
| First trial | G60 | 25.6 (3.8) | 25.6 (5.2) | 29.7 (6.5) | 44.9 (8.7) | 20.7 (4.6) | 13.7 (3.6) |
|  | G20 | 57.4 (5.0) | 45.9 (12.1) | 24.9 (6.1) | 48.6 (10.5) | 18.3 (3.5) | 8.8 (2.5) |
|  | G5 | 65.9 (9.3) | 41.2 (7.1) | 17.4 (4.5) | 55.2 (4.5) | 29.8 (12.3) | 5.6 (1.2) |
| Last trial | G60 | 34.2 (5.4) | 15.0 (2.6) | 4.3 (1.9) | 43.0 (10.0) | 7.6 (1.7) | 6.4 (2.6) |
|  | G20 | 69.6 (9.0) | 25.1 (7.6) | 11.9 (7.0) | 64.6 (10.9) | 25.5 (6.2) | 6.6 (2.1) |
|  | G5 | 70.5 (10.6) | 16.0 (6.3) | 6.0 (1.9) | 56.4 (7.2) | 8.7 (2.7) | 11.1 (5.4) |

**Figure 2.**
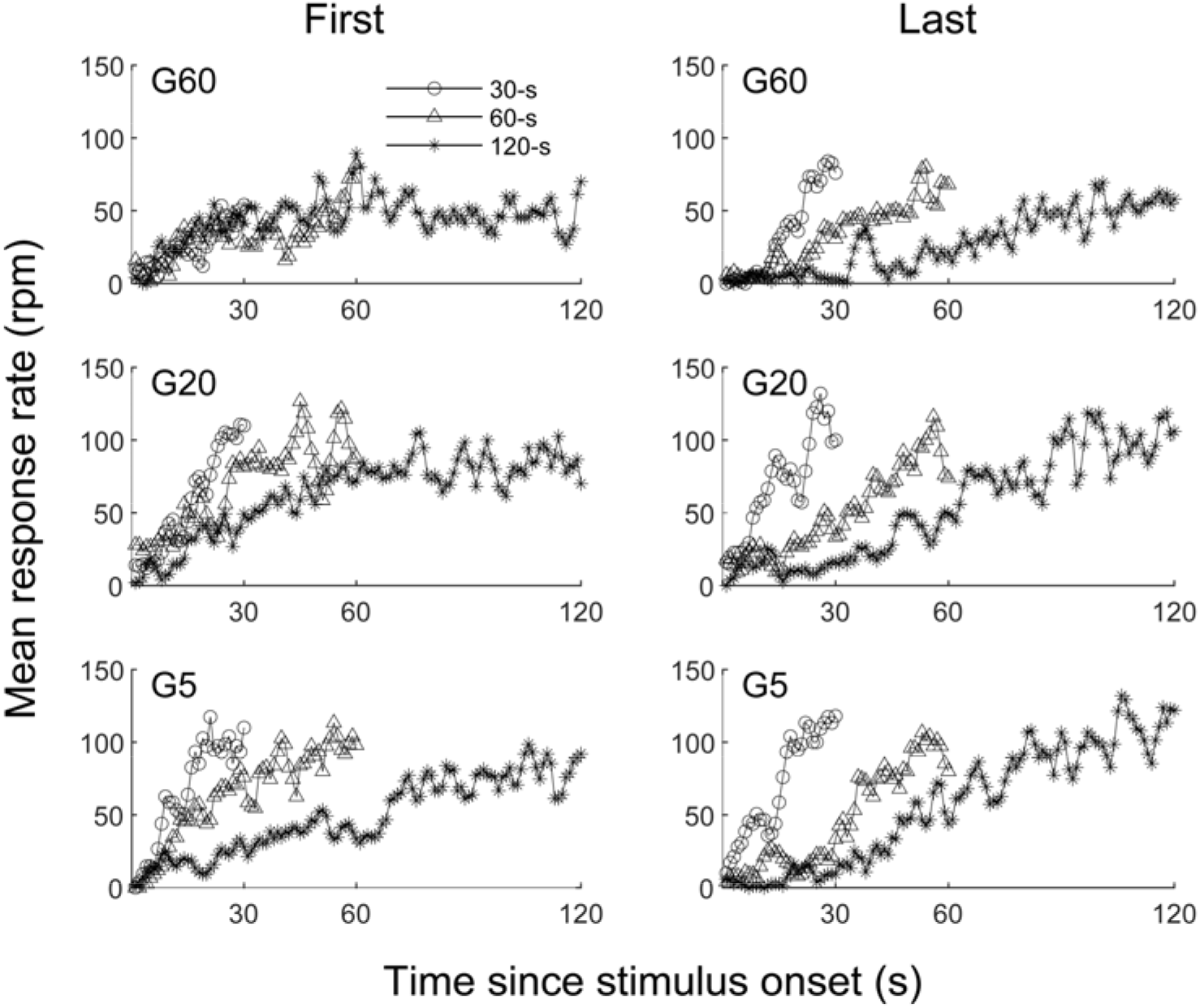
Mean response rate during the FI 30, FI 60, and FI 120 s as a function of time since stimulus onset in first (left column) and last (right column) trials at the end of blocked training for groups G60 (upper panels), G20 (middle panels), and G5 (lower panels).

Finally, Figure 3 shows mean response rate as a function of time from stimulus onset for groups G60 (top row), G20 (middle row), and G5 (bottom row) in the first (left column) and last (right column) trials of intermixed training. A repeated-measures ANOVA with interval (30-vs. 60-vs. 120-s FIs) and trial (first vs. last) as within-subject factors revealed a significant effect of interval for group G60 (*F*_*2,8*_ = 31.02, *p* < 0.001, η^²^_p_ = 0.889), for group G20 (*F*_*2,8*_ = 22.45, *p* < 0.001, η^²^_p_ = 0.849), and for group G5 (*F*_*2,8*_ = 49.61, *p* < 0.001, η^²^_p_ = 0.925). There were no significant effects of trial or interactions. These results suggest that rats from all groups similarly discriminated across the three FIs in the first and last trials of the final sessions of training.

**Figure 3.**
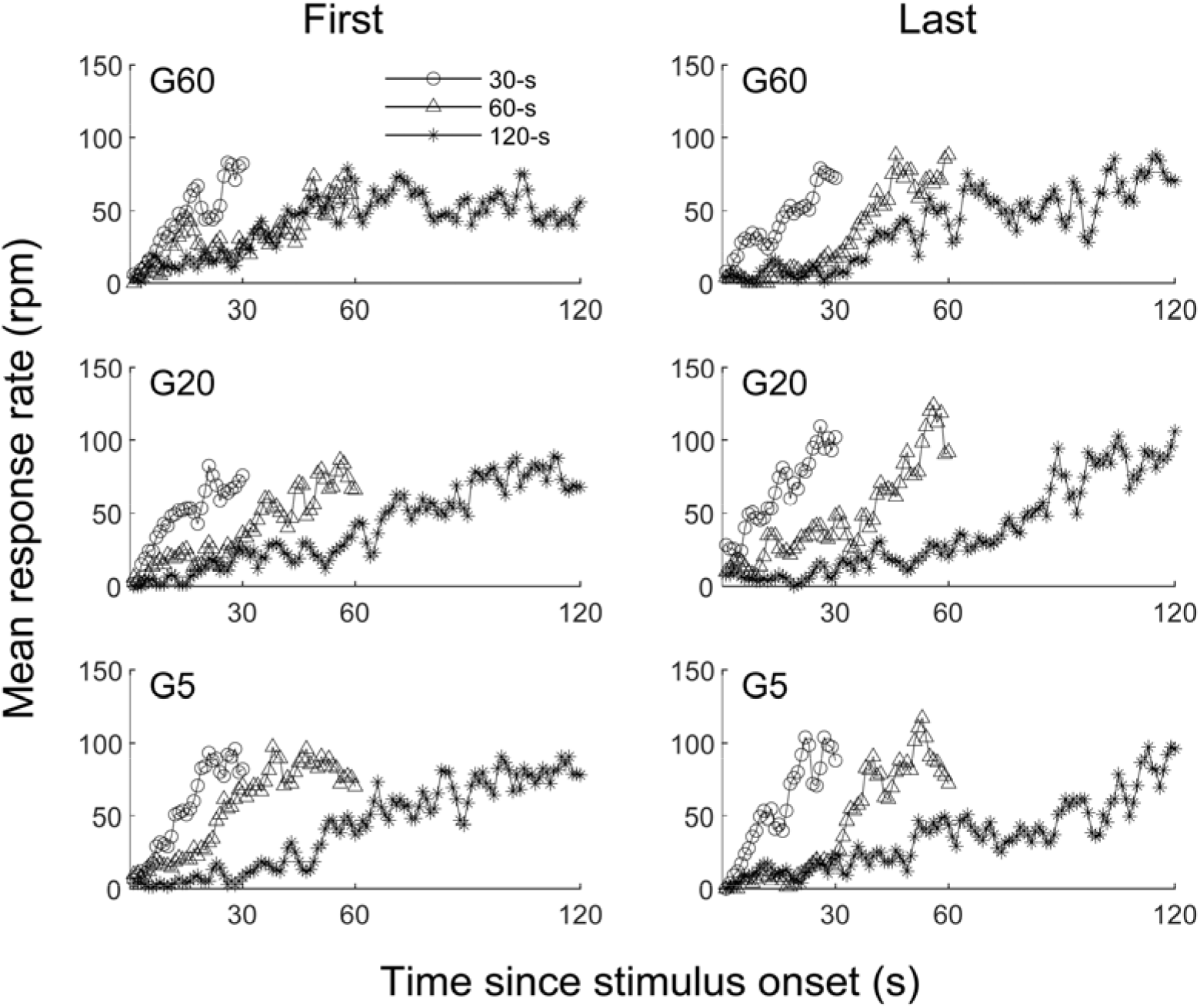
Mean response rate during the FI 30, FI 60, and FI 120 s as a function of time since stimulus onset in first (left column) and last (right column) trials at the end of intermixed training for groups G60 (upper panels), G20 (middle panels), and G5 (lower panels).

### 2.3 Discussion

In previous studies, we have shown that when FIs signaled by different stimuli are trained in a blocked design, stimulus control does not emerge (Caetano, Guilhardi, & Church, 2007, 2012). In Experiment I, rats were trained with three FIs signaled by three different stimuli in blocks of 60 consecutive trials (group G60), 20 consecutive trials (group G20) or 5 consecutive trials (group G5). The goal was to gradually decrease temporal regularity by adjusting block size, and by doing so, possibly identifying the point from which rats switch from using temporal regularity to using the stimuli as predictors to the FIs.

By the end of blocked training, animals in the group of greater temporal regularity (G60) did not discriminate across the different FIs early in each session, suggesting that the visual stimuli did not work as cues to the time of reinforcement in each trial. The animals clearly did not ignore the stimuli because response rate increased when any stimulus was turned on (as opposed to little to no responding during the ITI). However, the animals did not differentially use the stimulus to predict when food would be available, which led to overlapping gradients. In contrast, rats in groups G20 and G5 showed differential responding to each stimulus (i.e., stimulus-controlled performance) early in each session at the end of blocked training. Importantly, note that the overall number of trials of each FI trained during the experiment is the same across groups; the only difference is the order in which they were arranged.

When presented with intermixed trials, all rats showed stimulus-controlled performance, regardless of group membership. In this condition, the only possible way to correctly predict the time of food availability is to use the discriminative stimulus, while in the blocked condition the temporal regularity (i.e., time to reinforcement in the previous trials) can also be used as a cue to the time of food prime in the current trial. The fact that rats in group G60 were able to respond differentially to each stimulus when they were presented intermixed within the session rules out the possibility that those rats were simply unable to discriminate across the stimuli in the first place. The differences in performance between blocked and intermixed training for group G60 should then be a factor of the arrangement of the trials trained.

Overall, these results suggest that with decreasing temporal regularity, rats increasingly rely on the stimulus as a source of information about the time to reinforcement in each trial. Although these results agree with previous findings in humans using similar manipulations (Labliuk et al., 2015), it is desirable that future replications with larger sample sizes are carried out, as the number of rats in each group in this study was relatively small.

## 3. Experiment II

It is possible that, by reducing the number of signaled FIs, and therefore the number of associations that need to be learned, the stimuli could acquire control over performance even when trained in a blocked design. To test this hypothesis, in Experiment II rats were trained with only two signaled FIs, first in a blocked design and then with within-session intermixed trials.

### 3.1 Method

#### 3.1.1 Animals

Twelve male Wistar rats (purchased from Federal University of São Paulo, São Paulo - SP) arrived in the lab at 33 days of age. The conditions of maintenance and initial treatment of the animals were the same as described in Experiment I, except for the daily handling before the beginning of training, which lasted 32 days in this experiment. Training began one week after food restriction when they were 62 days old. All training sessions occurred in the morning.

#### 3.1.2 Apparatus

The same apparatus described in Experiment I was used in Experiment II, except for the discriminative stimuli used during training. Two visual stimuli were used during training: one was a solid light, referred to as “light”, which was located above the left lever; the other stimulus was a flickering light, referred to as “flicker”, located at the top of the wall opposite to the food cup.

#### 3.1.3 Procedure

Rats were trained on a multiple-fixed-interval procedure with two intervals (15 or 60 s) signaled by two different visual stimuli (light or flicker) for 36 daily sessions (Monday through Saturday). The assignment of visual stimuli to intervals was counterbalanced across rats. Experimental sessions consisted of 60 trials or 60 minutes, whichever came first. In each trial, food was primed either 15 or 60 seconds after stimulus onset. The first head entry into the food cup after the food was primed delivered the sucrose pellet, terminated the stimulus, and started a 20-s period with no stimulus (ITI). Head entries made prior to the time of food prime were recorded but had no effect.

##### 3.1.3.1 Phase 1 – Blocked training

In the first phase (30 sessions), animals were trained in blocks of 60 consecutive trials of one signaled FI per session, alternating every session (i.e., one FI per day), similar to arrangement described for group G60 in Experiment I.

##### 3.1.3.2 Phase 2 – Intermixed training

In the second phase (6 sessions), each session had 60 trials in which both signaled FIs were trained (30 trials of each). Each set of 20 trials was composed of 10 trials of each FI. A random permutation of those 20 trials defined the order in which they were trained. This procedure was repeated for the next 40 trials (in blocks of 20) and was done independently for each session and rat.

#### 3.1.4 Data analysis

As in Experiment I, temporal gradients were first averaged across the last three sessions of each FI in blocked and intermixed training. The dependent variable and percentage of sessions used for the statistical comparisons to describe performance of rats in Experiment II were the same as in Experiment I.

In order to make it possible to directly compare results across experiments and to keep the criteria for the individual analyses constant across experiments, mean response rate during the *comparable periods* (i.e., first 15 seconds) of each gradient was compared between FIs (15 vs. 60 s), phases (blocked vs. intermixed), and trials within sessions (first vs. last) in a repeated-measures ANOVA. Where applicable, partial eta squared or Cohen’s *d* was used to describe effect sizes. Post hoc Bonferroni corrections were used where appropriate. A test of normality (Shapiro-Wilk) was used to verify the assumption normality before all inferential analysis. All analysis revealed no deviation from normality. In order to show the behavioral adaptation to the FI schedule operating in each session during blocked training (data shown in Figure 5), we calculated the difference in performance between the 15 and 60-s gradients (i.e., subtracted the 15-s gradient from the 60-s gradient) trial by trial for each rat, then averaged across bins (i.e., we had one mean difference score per trial, session, and rat), then averaged across sessions and finally across rats.

Two rats did not discriminate between FIs during intermixed training and were excluded from the analyses. For both rats, the difference between mean response rates in the 15- and 60-s gradients was below 2.5 SEM (same criterion used in Experiment I).

### 3.2 Results

During blocked training, rats completed all 60 FI-15 and about 40 out of the 60 FI-60 trials during the 60-minute sessions. During intermixed training, all rats completed the 60 trials within the 60-minute sessions.

Figure 4A shows the temporal gradients for the 15- and 60-s FIs averaged across all trials at the end of the blocked and intermixed training. After training in both phases, rats discriminated between the two FIs, which is illustrated by the earlier rise in response rate for the 15-s interval compared to the 60-s interval in both panels. Mean rate (and standard error) for the comparable periods of the 15- and 60-s gradients during blocked training were 73.7 (3.9) responses per minute (rpm) and 19.4 (2.7) rpm, respectively; and 76.2 (5.1) rpm and 35.2 (4.6) rpm for the two gradients during intermixed training. A repeated-measures ANOVA with interval (15 vs. 60 s) and training (blocked vs. intermixed) as factors revealed a significant effect of interval (*F*_*1,9*_ = 207.76; *p* < 0.001, η^²^_p_ = 0.958), of training (*F*_*1,9*_ = 8.59; *p* = 0.017, η^²^_p_= 0.488), and a significant interval x training interaction (*F*_*1,9*_ = 15.28; *p* = 0.004, η^²^_p_ = 0.629).

**Figure 4.**
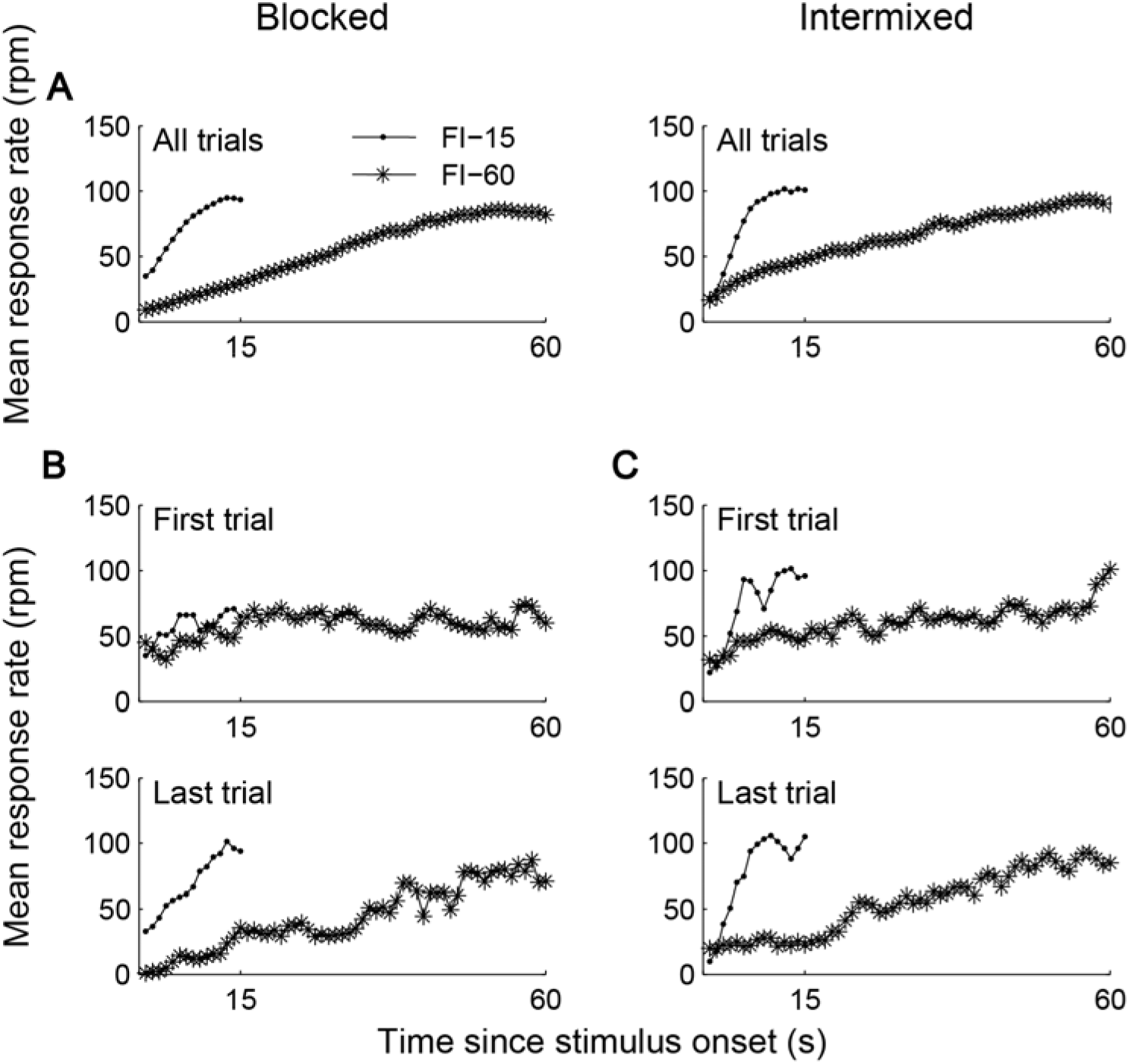
Mean response rate during the FI 15 s and FI 60 s as a function of time since stimulus onset. A) Overall mean across all trials within-sessions at the end of blocked (left panel) and intermixed (right panel) training. B) Mean response rate in the first (upper panel) and last (bottom panel) trials at the end of blocked training. C) Mean response rate in the first (upper panel) and last (bottom panel) trials at the end of intermixed training.

To further investigate the source of the significant interaction observed, a post hoc comparison between blocked and intermixed training both for the 15 and 60-s gradients was performed. Post hoc comparisons showed no significant effect of training for the 15-s gradient (*t*_*9*_ = 0.64, *p* = 0.539) but a significant effect of training for the 60-s gradient (*t*_*9*_ = 4.95, *p* < 0.001, Cohen’s *d* = 1.564). There was also a significant effect of interval both in blocked (*t*_*9*_ = 12.57, *p* < 0.001, Cohen’s *d* = 3.974) and in intermixed training (*t*_*9*_ = 13.67, *p* < 0.001, Cohen’s *d* = 4.324).

Although the temporal gradients in Figure 4A suggest similar performance after intermixed and blocked training, individual-trial analyses show differences in temporal performance between the two training types. Figure 4B shows performance in the first and last trials within sessions during blocked and intermixed training. At the end of blocked training (left panels), the two temporal gradients superposed early in the session and separated later in the session. Mean rate (and standard error) for the first 15 s of each gradient in the first trial were 58.0 (6.4) rpm and 46.2 (5.5) rpm for the 15 and 60-s gradients, respectively; and 65.8 (4.2) rpm and 14.2 (3.6) rpm in the last trial. A repeated measures ANOVA with two within-subject factors, interval (15 vs. 60 s) and trial (first vs. last), showed a significant effect of interval (*F*_*1,9*_= 33.68, *p* < 0.001, η^²^_p_ = 0.789), of trial (*F*_*1,9*_ = 5.66, *p* = 0.041, η^²^_p_ = 0.386), and a significant interval x trial interaction (*F*_*1,9*_ = 13.43, *p* = 0.005, η^²^_p_ = 0.599). This interaction expresses the adjustment in performance observed across trials.

Post hoc comparisons showed no significant difference between the 15 and 60-s gradients in the first trial (*t*_*9*_= 1.32, *p* = 0.221), but revealed a significant difference between the two gradients in the last trial (*t*_*9*_ = 8.33, *p* < 0.001, Cohen’s *d* = 2.633). Also, post hoc comparisons showed no significant difference between first and last trials for the 15-s gradient (*t*_*9*_ = 1.07, *p* = 0.313), but a significant difference between trials for the 60-s gradient (*t*_*9*_ = 4.21, *p* = 0.002, Cohen’s *d* = 1.332).

During intermixed training, on the other hand, temporal performance was already a function of signaled FI from the first trial (Figure 4C). Mean rate (and standard error) for the first 15 s of each gradient in the first trial were 74.4 (7.3) rpm and 60.2 (4.3) rpm for the 15 and 60-s gradients, respectively; and 76.7 (6.2) rpm and 54.5 (7.0) rpm in the last trial. A repeated measures ANOVA revealed a significant effect of interval (*F*_*1,9*_ = 18.27, *p* = 0.002, η^²^_p_ = 0.670), no effect of trial (*F*_*1,9*_ = 0.08, *p* = 0.781), and no interval x trial interaction (*F*_*1,9*_ = 0.31, *p* = 0.593).

In order to further describe the within-session adjustment in performance observed during blocked training, we calculated a mean difference score between the 15- and 60-s gradients trial-by-trial during the last three sessions of blocked training (Figure 5). Low scores suggest superposed gradients, and the higher the score the larger the difference between gradients. As already shown in Figure 4B, the two temporal gradients superposed early in the sessions (low difference scores for trials 1 and 2) but quickly separated as a function of trials. By the fifth trial, the difference between gradients reached a plateau and remained approximately constant until the end of the session.

**Figure 5.**
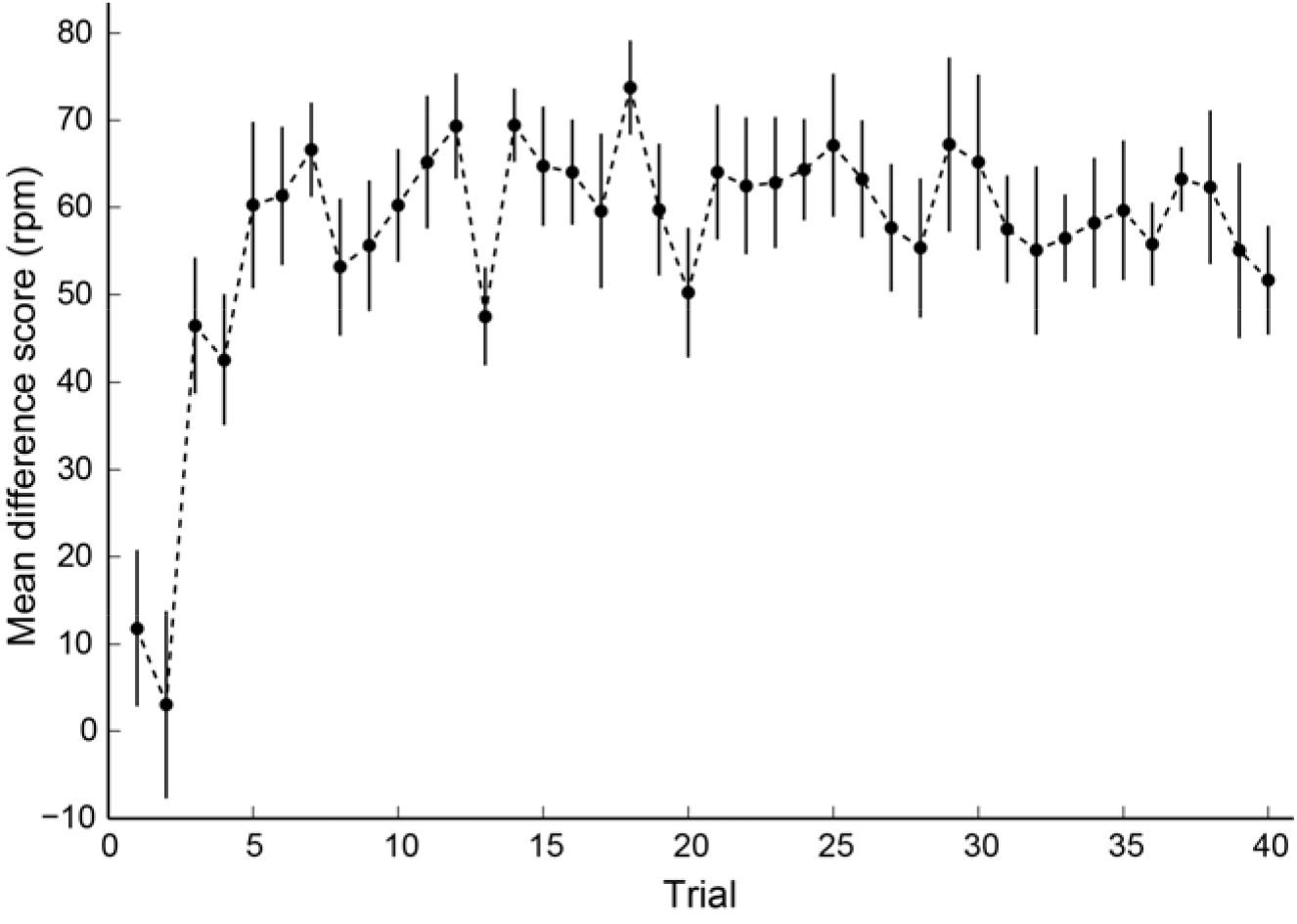
Mean difference score between the FI 15 and FI 60 s gradients as a function of trials during blocked training.

### 3.3 Discussion

In Experiment I, we manipulated temporal regularity by controlling the number of consecutive trials within blocks during blocked training, and assessed its effect on the control of temporal performance. From these results, we hypothesized that by training a reduced number of FIs (and thus by requiring fewer associations between stimuli and intervals to be formed) the stimuli could acquire control over temporal behavior even when trained in blocks. Therefore, in Experiment II we trained rats with only two signaled FIs. Contrary to our initial hypothesis, after blocked training the stimuli did not acquire control over behavior. Instead, animals showed a rapid behavioral adjustment at the beginning of each blocked session to adapt to the interval trained in each day (Figure 5). This was illustrated by the overlapping temporal gradients in the first trial, and separate gradients in the last trial of each session – similarly to what was observed in Experiment I for rats trained in group G60, in the blocked condition. These results suggest that whenever there are temporal regularities in the task which could signal when an event will occur, stimulus control over temporal performance does not emerge, even in the situation where there is a small number of associations between stimuli and FIs to be learned. As noted in Experiment I, these results do not mean that rats simply ignored the stimuli, but rather that they did not use them as different cues to different times to reinforcement.

When trained with intermixed trials, on the other hand, temporal gradients were clearly distinct starting from the first trial. Similar to Experiment I, this control condition ensured that any failure in developing stimulus-controlled performance was not due to an inability to discriminate between stimuli.

In summary, Experiment II suggests that when there is a temporal regularity across trials, the visual stimuli do not acquire control over temporal performance even in the situation where there is a small number of associations between stimuli and FIs to be learned. However, an open question is how strong such temporal regularity must be in order to control behavior.

## 4. General discussion

The results herein described suggest that temporal regularity strongly controls temporal performance. Whenever possible, rats quickly adjust performance based on previous adjacent trials rather than show stimulus-controlled performance. We were able to investigate the emergence of stimulus control by training different FIs either in blocks of several consecutive trials – thus controlling the temporal regularity – or with intermixed trials within sessions, as previously reported (Caetano et al., 2007, 2012). As rats adjusted their performances within very few trials, analyses based on averages across trials would not show differences between blocked and intermixed designs. Indeed, when trial averages were compared between blocked and intermixed training in Experiment I (Figure 1) and in Experiment II (Figure 4, top panels), temporal gradients were indistinguishable across conditions, suggesting similar stimulus and interval discrimination. This illustrates the importance of investigating individual trials, as some interesting phenomena can be hidden by averages (Church, Meck, & Gibbon, 1994; Gallistel, Fairhurst, & Balsam, 2004).

It is plausible that there is a hierarchy of simplicity in the strategy adopted by rats in multiple temporal discriminations, as previously suggested (Caetano et al., 2012). That is, it may be easier to attend to the temporal regularities in the task than to learn relationships between stimuli and FIs. Alternatively, our findings are compatible with the idea of a competition for associative strength between the visual stimuli and the temporal cues. This phenomenon, known as *overshadowing*, is observed when a more salient stimulus obscures a less salient stimulus. The more salient stimulus acquires more strength of association and control over behavior. Our results suggest that temporal regularity overcomes the stimuli in the control for temporal performance, as previously suggested by Marshall & Kirkpatrick (2015).

The phenomenon of overshadowing was first described by Pavlov in 1927. In his studies with dogs, Pavlov noted that when two or more cues signal the same event, they compete for the control of the response, and depending on the characteristics of the cues (e.g., salience) one will end up eliciting more responses than the other (Pavlov, 1927; Rescorla, 1988). Since Pavlov’s series of experiments, overshadowing has been extensively observed and described in other experiments with classical and also operant conditioning.

While some studies investigated the effect of the duration of simultaneously presented stimuli in the competition for associative strength (Bonardi, Mondragón, Brilot, & Jennings, 2015; Dómhnall, Eduardo, Esther, & Charlotte, 2011; D. J. Jennings, Alonso, Mondragon, Franssen, & Bonardi, 2013; Domhnall J. Jennings, Bonardi, & Kirkpatrick, 2007; McMillan & Roberts, 2010), others reported the effect of the training arrangements involving multiple FIs on the development of stimulus control over temporal performance (Caetano et al., 2007, 2012; Guilhardi et al., 2010; Labliuk et al., 2015). Along these lines, decreasing block size in Experiment I could be interpreted as leading to a corresponding decrease in salience in temporal regularity, which resulted in an increase in stimulus-controlled performance.

In humans, Guilhardi and colleagues (2010) used a modified peak procedure task to test whether the discriminability across stimuli played a role in the development of stimulus control over temporal performance. Participants performed a shooting task in which a moving target traveled from left to right on the computer screen. Shots could be given at the center of the screen. The goal was to maximize the number of hits and avoid missing the target (i.e., avoid shooting too early or after it passed the center of the screen). In regular trials, the target and its trajectory could be seen. However, during test trials, both the target and its trajectory were masked, so participants had to estimate the time taken for the target to reach the center of the screen to shoot at the appropriate time. Three background colors were paired with three different target speeds. A lighter, darker, and intermediate green (manipulation of brightness) were associated with a faster, slower and intermediate target speed. Trials were first presented in blocks (60 consecutive trials of each type), followed by 45 test trials (with occluded target and trajectory) randomly presented to test whether or not participants associated the background colors to their target speeds. Finally, all trials were then trained intermixed and a new set of test trials followed to make sure participants could learn those associations. Results showed that, after training in blocks, participants did not associate background colors to target speeds. That is, performance on the three trial types was indistinguishable when the target could not be seen. After trained with intermixed trials, on the other hand, participants discriminated target speeds based on background colors even when its trajectory was masked.

Using the same shooting task, Labliuk et al. (2015) showed that the number of consecutive trials within blocks played a role on whether temporal performance became a function of the background colors. The more trials per block, the less likely performance were controlled by the stimuli. Block size varied from 1 (i.e., intermixed) to 60 (blocked), with intermediate blocks containing 10, 20 and 30 trials. Importantly, the amount of training of each signaled FI was held constant across all conditions; the only difference was the training arrangement (block size).

The effect of the order in which the different FIs are trained – in blocks of consecutive trials or intermixed across trials – cannot be predicted by current models of timing. Scalar Expectancy Theory – SET (Gibbon, 1977; Gibbon, Church & Meck, 1984), for example, proposes an internal clock mechanism composed of a pacemaker, which emits pulses when triggered by a relevant stimulus; an accumulator, which stores those pulses; and a comparator which compares the number of pulses stored in the comparator to an exemplar randomly selected from a reference memory. This reference memory is formed by pulses accumulated in previous reinforced trials. The model assumes that distinct reference memories are formed when different stimuli signal different FIs. Therefore, the signaling stimulus in each trial indicates from which reference memory the exemplar must be drawn. The order in which different signaled FIs is trained is irrelevant to the formation of such reference memories. That is, whether different FIs are trained in a blocked or intermixed design, the resulting memories formed are identical, which leads to an incapacity of the model in reproducing the results herein reported.

A behavioral alternative model to SET, called Learning-to-time – LeT (Machado, 1997), similarly fails to account for those results. LeT assumes that a series of behavioral states get differentially linked to a terminal response as training in fixed interval procedures progresses. Similar to the pulses in SET, a vector of the association strength between each behavioral state and the terminal response represents the memory for the timed interval. Different from SET, this singular memory (as opposed to a collection of pulses from previous experiences in SET) is reshaped with every reinforced or non-reinforced trial. However, similar to SET, the model assumes that different strength vectors are formed as different stimuli signal distinct FIs, which also results in an inability of the model to account for the effect of order of training described in the present set of experiments.

The effect of the order in which FIs are trained on temporal performance seem robust, as they have been replicated in rats (Caetano et al., 2007, 2012) and humans (Guilhardi et al., 2010; Labliuk et al., 2015). It is possible that current models of timing and conditioning can be adapted to take into account the training arrangement in their predictions.

## Acknowledgements

Support for this study was provided to EBN by UFABC and process #2014/22918-2, São Paulo Research Foundation (FAPESP), to MSC by the National Counsel of Technological and Scientific Development (CNPq), Grant #485272/2013-0, and to MBR by the National Counsel of Technological and Scientific Development (CNPq), Grant #430993/2016-1. MSC is affiliated to Instituto Nacional de Ciência e Tecnologia sobre Comportamento, Cognição e Ensino, with support from the Brazilian National Research Council (CNPq, Grant # 465686/2014-1), the Coordination of Superior Level Staff Improvement (CAPES, Grant # 88887.136407/2017-00), and the São Paulo Research Foundation (Grant # 2014/50909-8). The authors would like to thank Armando Machado, Russ Church and the members of the Timing and Cognition Laboratory at UFABC (http://neuro.ufabc.edu.br/timing/) for useful discussions and suggestions on this study.

